# Bioconversion of *p*-coumaric acid to *cis,cis*-muconic acid using an engineered *A. baylyi* ADP1 - *E. coli* co-culture

**DOI:** 10.64898/2026.03.05.709578

**Authors:** Suchismita Maiti, T. Priyadharshini, Lars M. Blank, Guhan Jayaraman

**Affiliations:** Department of Biotechnology, Bhupat and Jyoti Mehta School of Biosciences, Indian Institute of Technology Madras, Chennai, Tamil Nadu, 600036, India; Institute of Applied Microbiology, RWTH Aachen University, Worringerweg 1, 52074, Aachen, Germany

**Keywords:** *A. baylyi* ADP1, *cis,cis*-muconic acid, *p*-coumaric acid, β-ketoadipate pathway, co-culture system, dual-substrate feeding, lignin valorization

## Abstract

Lignin-derived aromatics are abundant in depolymerized lignin but remain remain untilized as carbon sources for commercial production of bulk chemicals. Among these aromatics, *p*-coumaric acid can be funnelled through the β-ketoadipate pathway toward *cis,cis*-muconic acid (ccMA), a precursor of bio-based adipic and terephthalic acids. However, efficient ccMA production by *Acinetobacter baylyi* ADP1 is constrained by toxicity of catechol (the immediate precursor of ccMA), inefficient channelling of protocatechuate (PCA) metabolism towards ccMA production, and absence of PCA decarboxylase for converting PCA to catechol. Therefore, in this study, we engineered a modular co-culture system, combining engineered strains of *A. baylyi* and *E. coli*, for ccMA production from synthetic *p*-coumaric acid. Deletion of *catB* and *catC* genes and overexpression of *catA* in *A. baylyi* GJS_*catA* strain enabled near-stoichiometric conversion of catechol to ccMA (∼90% carbon yield) with titres up to 56.4 mM (∼ 8 g/L) under controlled fed-batch feeding. The strain was further engineered (*A. baylyi* GJS2_*catA*) to convert *p*-coumaric acid to PCA. Due to the inactivity of heterologous PCA decarboxylase (*aroY* gene) in *A. baylyi*, this gene was incorporated in *E. coli* where it exhibited activity through PCA to catechol conversion. Upon its production by *E.coli_aroY* in the co-culture, catechol is instantaneously converted to ccMA by *A. baylyi* GJS2_*catA* strain. In a two-step process, 22 mM *p*-coumaric acid was initially converted to 20.6 mM PCA (*A. baylyi* GJS2_*catA*), which was further converted to catechol (*E.coli_aroY*) and finally to 18.55 mM ccMA (2.63 g L⁻¹) by *A. baylyi* GJS2_*catA*. This process was validated by the valorization of lignin-derived *p*-coumaric acid to ccMA. While the modular strategy developed in this study substantially improves ccMA titres, it also highlights the bottlenecks in *A. baylyi* metabolic pathway engineering for lignin valorization.

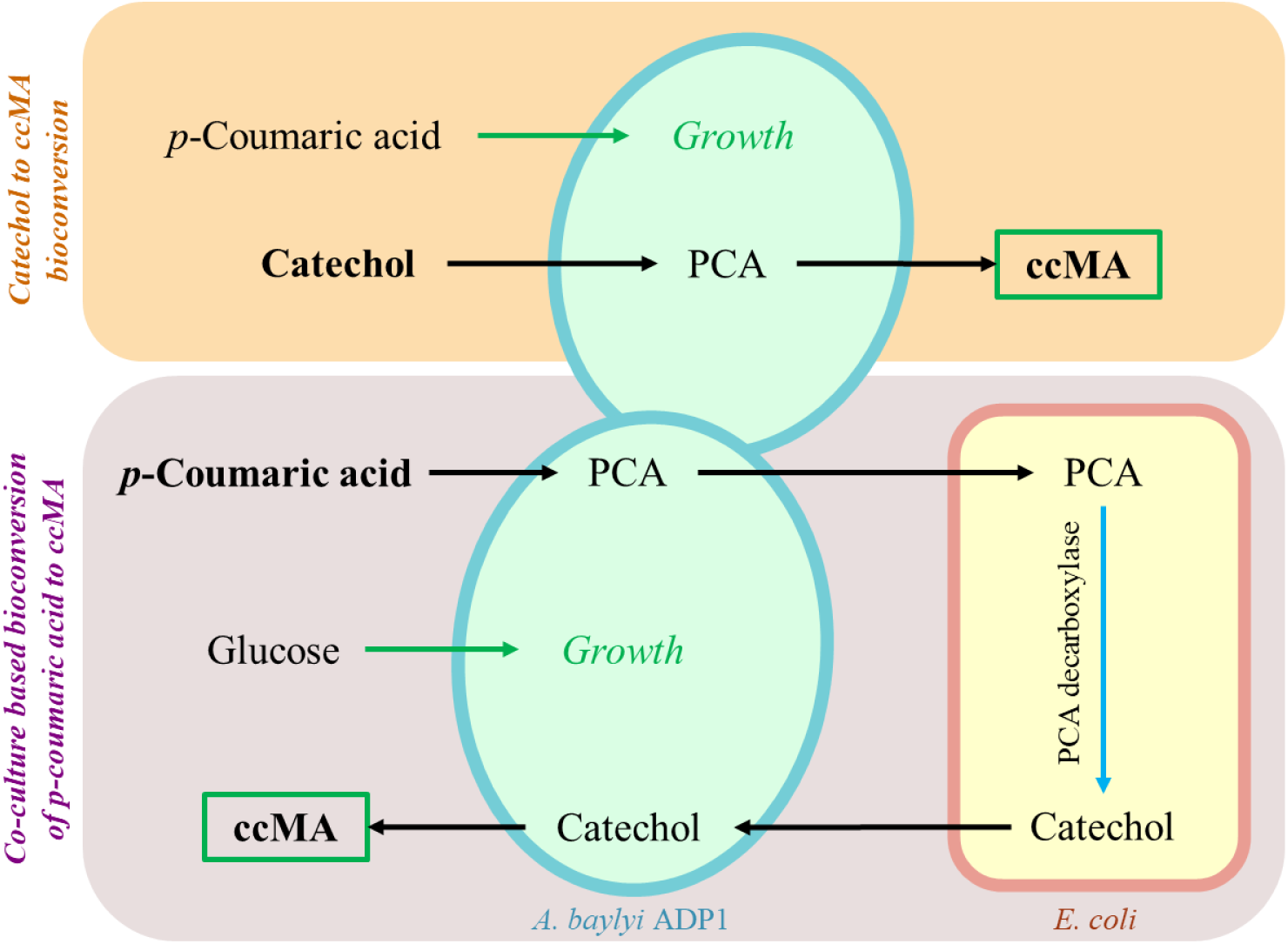

## 1. INTRODUCTION

Lignin-derived phenolic acids are an abundant but underused carbon source in lignocellulosic biorefineries. Among these, *p*-coumaric acid is especially attractive. It occurs at high levels in herbaceous and agricultural residues. When pretreated corncob and similar feedstocks are subjected to base-catalysed depolymerization, *p*-coumaric acid typically appears as the predominant monomer, with minor amounts of ferulic acid and other hydroxycinnamic acids (Sun et al., 2018; Sunkar & Bhukya, 2022). This makes *p*-coumaric acid a key substrate in the valorization of lignin-rich waste streams through the β-ketoadipate pathway, which is present in many bacteria, such as *Pseudomonas putida* and *Acinetobacter baylyi* ADP1. Converting such aromatics to *cis,cis*-muconic acid (ccMA) is particularly appealing because ccMA can be upgraded to adipic and terephthalic acids, which are central building blocks for high-performance polymers (Salvachúa et al., 2018).

The biosynthetic production of ccMA by bacterial fermentations has emerged as a sustainable and viable alternative to traditional petrochemical processes. Over the past decade, bacterial metabolic engineering has enabled considerable enhancements in ccMA titers, yields, and productivity, establishing it as a key biobased platform chemical with potential for industrial scale-up (Johnson et al., 2016; Salvachúa et al., 2018). Genetic engineering efforts have optimized two main biosynthetic routes for ccMA production through the key intermediate, PCA: channelling the sugars through the shikimate pathway or aromatics such as p-coumaric acid through the β-ketoadipate pathway to PCA, which can then be converted to ccMA either natively or with engineered modifications to improve flux (Fig. 1). Recent advances have demonstrated that engineering strains for efficient operation of both synthetic routes allows for flexible production from a variety of renewable feedstocks, including complex mixtures present in lignocellulosic hydrolysates (Table 1).

**Figure 1.**
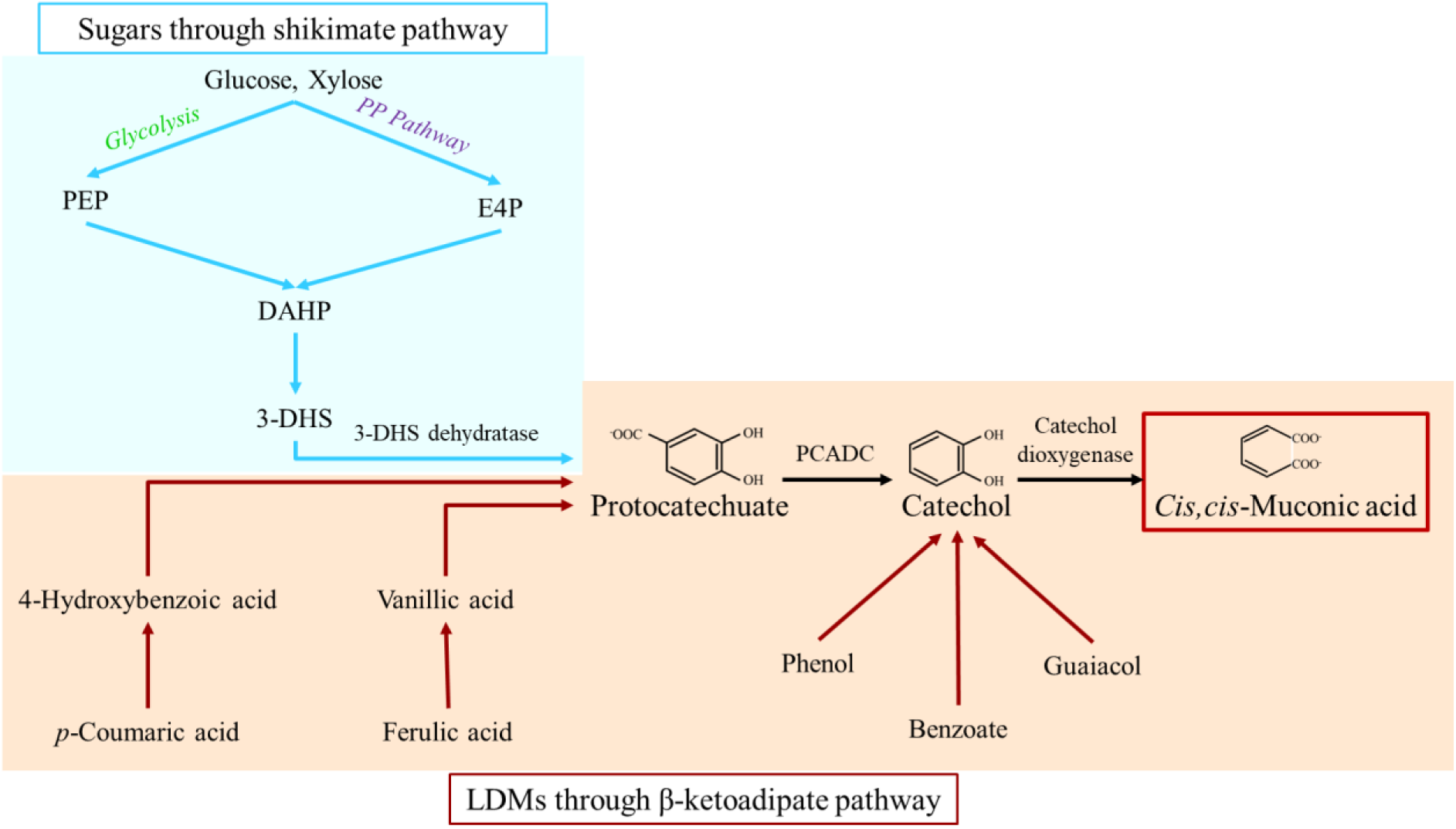
Schematic showing the shikimate pathway and β-ketoadipate pathway for ccMA production. The abbreviations here are as follows: PP Pathway - Pentose Phosphate Pathway, PEP - Phosphoenol pyruvate, E4P - D-erythrose 4-phosphate, DAHP - 3-deoxy-D-arabino-heptulosonate-7-phosphate, 3-DHS - 3-dehydroshikimate.

**Table 1.** Different organisms producing *cis,cis*-muconic acid from different substrates.

| Type of substrates | Substrate | Engineered Bacteria | Titer (g/L) | Reference |
| --- | --- | --- | --- | --- |
| Synthetic substrates | Glucose | *Corynebacterium glutamicum* | 88.2 | Li et al., 2024 |
|  | Glucose and catechol | *Corynebacterium glutamicum* | 85 | Becker et al., 2018 |
|  | Catechol | *Pseudomonas putida* KT2440 | 64.2 | Kohlstedt et al., 2018 |
|  | Glucose and *p*-coumaric acid | *Pseudomonas putida* KT2440 | 50 | Salvachúa et al., 2018 |
|  | Glucose and Xylose (corn stover hydrolysate) | *Pseudomonas putida* KT2440 | 47.2 | Mokwatlo et al., 2024 |
|  | Glucose and Xylose | *Pseudomonas putida* KT2440 | 33.7 | Ling et al., 2022 |
|  | Glucose | *Pseudomonas putida* KT2440 | 22 | Bentley et al., 2020 |
|  | Glucose | *Saccharomyces cerevisiae* | 20.8 | Wang et al., 2020 |
|  | Glucose and *p*-coumaric acid | *Pseudomonas putida* KT2440 | 15.6 | Johnson et al., 2016 |
|  | Gluconate, *p*-Coumaric acid and Ferulic acid | *Acinetobacter baylyi* ADP1 | 0.48 | Liu et al., 2025 |
|  | *p*-Coumaric acid and catechol | *Acinetobacter baylyi* ADP1 | 8 | This study |
|  | Glucose and *p*-Coumaric acid | Co-culture of *A. baylyi* ADP1 and *E.coli* | 2.63 | This study |
| Depolymerized lignin | Pine lignin (depolymerized and concentrated, containing Catechol, phenol and cresol) | *Pseudomonas putida* KT2440 | 13 | Kohlstedt et al., 2018 |
|  | Cornstover (base-catalyzed containing *p*-Coumaric acid and ferulic acid) | *Pseudomonas putida* KT2440 | 3.70 | Salvachúa et al., 2018 |
|  | Pine lignin (Hydrothermal treatment, containing Catechol and phenol) | *Corynebacterium glutamicum* | 1.8 | Becker et al., 2018 |
|  | Kraft lignin (base catalyzed, containing Vanillin and guaiacol) | *Pseudomonas putida* KT2440 | 1.4 | Almqvist et al., 2021 |
|  | Cornstover lignin (treated with anthraquinone and NaOH, containing *p*-Coumaric acid, ferulic acid, acetate, glycolate) | *Pseudomonas putida* KT2440 | 0.7 | Vardon et al., 2015 |
|  | *p*-Coumaric acid extracted from depolymerized lignin (obtained alkali-treated lignin-rich residue of pre-treated, enzymatically hydrolysed corncob biomass) and catechol | *Acinetobacter baylyi* ADP1 | 1.2 | This study |
|  | Glucose and *p*-Coumaric acid extracted from depolymerized lignin | *Acinetobacter baylyi* ADP1 and *E.coli* | 0.24 | This study |

*Acinetobacter baylyi* ADP1 is a promising chassis for ccMA production, as it is naturally competent, shows high homologous recombination rates, and carries a well-characterized β-ketoadipate pathway (Draths & Frost, 1994). Unlike *Corynebacterium glutamicum* and *Pseudomonas putida*, *Acinetobacter baylyi* has not been sufficiently explored for ccMA production. A recent study by Liu et al. (2025) demonstrated successful engineering of *A. baylyi* ADP1 for ccMA production, achieving 83.4% molar yield from *p*-coumaric acid and ferulic acid by deleting regulatory genes (*catM, benR*) along with other pathway genes and overexpression of heterologous gallic acid decarboxylase (*Blastobotrys adeninivorans*). This study used gluconate as the primary substrate for growth. However, this approach required multiple gene deletions to overcome transcriptional repression of the protocatechuate (PCA) branch, which may compromise strain robustness and adaptability under industrial conditions. Also, the final ccMA titer was low (0.48 g/L) when compared to the other studies mentioned in Table 1.

In this study, *A. baylyi* Δ*IS* ADP1 obtained from Prof. Barrick’s lab (UT Austin), was validated for ccMA production. Deletion mutants (Δ*IS1236*) of these insertion sequences in ADP1 have demonstrated reduced mutation rates and improved transformation efficiency (Suárez et al., 2017). Thus, utilising the ADP1-ISx strain (named as *A. baylyi* Δ*IS* ADP1 in this study) for strain engineering would be advantageous for achieving long-term stable expression of heterologous genes for ccMA production. Initially, the strain was engineered for the production of ccMA from catechol in the presence of a secondary substrate for growth. This dual-substrate approach aims to decouple primary metabolism from product formation, potentially enhancing overall process efficiency while mitigating substrate toxicity effects and maintaining cellular viability throughout the bioconversion process.

Subsequently, we investigated the conversion of *p*-coumaric acid to ccMA via the key intermediates, PCA and catechol. Bioconversion of PCA to catechol requires the expression of a heterologous gene encoding protocatechuate carboxylase. Since this gene is missing in *A. baylyi* ADP1, the *aroY* gene from *Klebsiella pneumonieae* was expressed in the bacteria. The aroY enzyme was not found to be active in *A. baylyi* ADP1 upon expression, but was actively able to convert PCA to catechol in recombinant *E. coli* strain. This necessitated a co-culture process for the successful conversion of *p*-coumaric acid to ccMA.

In the final part of this study, we explored the bioconversion of depolymerized lignin, predominantly containing *p*-coumaric acid, to ccMA.

## 2. METHODS

### 2.1. Bacterial strain and growth conditions

The *A. baylyi ΔIS* ADP1 strain was provided by Prof. Jeffrey E. Barrick (U.Texas, Austin). Minimal Salt Medium (MSM) was used for all the experiments containing (L^-1^): K_2_HPO_4_ -3.88 g, NaH_2_PO_4_ - 1.63 g, (NH_4_)_2_SO_4_ - 2.00 g, MgCl_2_·6H_2_0 - 0.1 g, EDTA - 10 mg, ZnSO_4_·7H_2_O - 2 mg, CaCl_2_·2H_2_O - 1 mg, FeSO_4_·7H_2_O - 5 mg, Na_2_MoO_4_·2H_2_O - 0.2 mg, CuSO_4_·5H_2_O - 0.2 mg, CoCl_2_·6H_2_O - 0.4 mg, MnCl_2_·2H_2_O - 1 mg. Appropriate carbon sources were added in each growth study. All shake flask growth conditions were maintained at 275 rpm, pH 7, and 30 °C for *A. baylyi* ADP1 and 200 rpm and 37 °C for *E.coli* .

### 2.2. Genetic engineering of *A. baylyi* ADP1 and *E.coli*

The pBW162 plasmid (Addgene #140634) was used as the primary vector for gene expression in *A. baylyi* ADP1, in accordance with its prior use for mevalonate pathway genes (Arvay et al., 2021; Biggs et al., 2020). Gibson assembly protocols were used to construct plasmids mentioned in Table S1 (https://barricklab.org/twiki/bin/view/Lab/ProtocolsGibsonCloning).

Chromosomal gene knockouts in *A. baylyi* ADP1 were performed as described in the protocols given by Barrick’s Lab (Barricklab.org), using overlap PCR with *tdk/kan* cassette (Santala et al., 2014). For more details refer the link - https://barricklab.org/twiki/bin/view/Lab/ProtocolsAcinetobacterOverlapPCRTransformation. To construct a strain capable of accumulating ccMA from catechol, targeted gene deletions and overexpression modifications were implemented in the *A. baylyi* Δ*IS* ADP1 strain. The genes *catB* (ACIAD1446) and *catC* (ACIAD1447) were deleted. This strain was named the *A. baylyi* GJS strain. To enhance the bioconversion efficiency of catechol to ccMA, the *catA* gene (ACIAD1442) was subsequently cloned using Gibson-assembly into plasmid pBWB162 and overexpressed using an IPTG-inducible expression system. This strain was named the *A. baylyi* GJS_*catA* strain. To construct a strain capable of accumulating PCA from *p*-coumaric acid, the genes *pcaH* (ACIAD1711) and *pcaG* (ACIAD1712) were deleted in the *A. baylyi* GJS strain. Furthermore, the *acr1* gene (wax ester formation) was deleted, and the strain named as *A. baylyi* GJS2. Catechol decarboxylase (*catA* gene) was also overexpressed in this strain. Wild-type *aroY* gene (PCA decarboxylase) was synthesized by Twist Bioscience Pvt. Ltd. and cloned into *E. coli* for PCA to catechol bioconversion. This list of all the strains generated in this study is mentioned in Table S2.

### 2.3. Shake flask studies for *cis, cis* – Muconic acid production from catechol

Since A. baylyi ADP1 grows on different carbon substrates, we investigated the effect of an independent growth-substrate on the bioconversion of catechol to ccMA. Three different growth-substrates - glucose, acetate and *p*-coumaric acid, were evaluated in independent shake-flask experiments. Duplicate cultures of *A. baylyi* GJS_*catA* strain were grown in 250 mL Erlenmeyer flasks containing 25 mL of mineral salt medium (MSM) at 30 °C and 250 rpm. The initial concentration of each growth-substrate in the flask was kept as follows: 30 mM glucose, 60 mM sodium acetate, or 20 mM *p-*coumaric acid. The growth-substrates were further added in a step-wise manner until the culture reached the late-exponential to early-stationary phase, as indicated by an optical density (OD_600_) of 6 -10. Upon reaching the target OD_600_ range, catechol was added at 3 h intervals, and brought to a concentration of 10 mM. Samples were collected periodically to monitor cell density and to quantify catechol and ccMA by HPLC.

### 2.4. Bioreactor studies for *cis, cis* – Muconic acid production from catechol

Bioreactor cultivations (duplicate experiments) were performed with dual-substrates in a 1.2 L bioreactor (Bioengineering AG, Switzerland) at 30 °C, pH 7-7.5 and 300 rpm, to evaluate the ccMA production. The *A. baylyi* GJS_*catA* strain was inoculated in mineral salt medium with an initial OD_600_ of 0.5. Dissolved oxygen (DO) was regulated at 20% by cascade control of airflow rate and agitation speed. The primary carbon source, *p-*coumaric acid, was initially kept at 20 mM and supplemented from a stock of 0.5 M *p-*coumaric acid whenever it was depleted in the fermentation medium. Catechol feed was initiated at late exponential phase (OD_600_ 9-10) at a constant rate of ∼ 3 millimoles.h⁻¹ from a 2 M stock solution, in order to maintain a 2.5 mM catechol concentration within the bioreactor. Along with the catechol feed, 5 millimoles *p-*coumaric acid was added every 6 hours through a separate pump. Samples were collected every 3 h and analyzed for growth (OD_600_) and for substrate and product concentrations by HPLC.

### 2.5. Co-culture studies for bioconversion of *p-*coumaric acid to ccMA

These experiments were done in duplicates in 250 ml shake flasks with 25 ml culture volume. *A. baylyi* GJS2_*catA* was cultivated in mineral salt media initially containing 5 mM glucose and 3 mM *p-*coumaric acid. Glucose was further supplemented upon depletion, and *p-*coumaric acid was supplemented upon conversion to PCA. HPLC analysis was done during the shake-flask experiments to monitor the concentration of these 2 carbon sources. As soon as 20 mM PCA was accumulated, 10 OD_600_ of *E.coli*_*aroY* was added to the culture for the whole-cell bioconversion of PCA to catechol.

### 2.6. Extraction of *p-*Coumaric acid from Corncob

Alkali pretreated lignin from the lignin-rich fraction (LRF) of corncob was obtained as mentioned in Sivapuratharasan et al., 2025. To obtain *p-*coumaric acid, alkali pretreatment of LRF, at 15% solid loading, was performed using 1 M NaOH for 24 hours at room temperature. The alkali-pretreated liquor was centrifuged, and *p*-coumaric acid was precipitated along with other aromatics by acidification of the pretreated liquor to pH 2, using concentrated HCl. The acidified solid precipitate was mixed with ethyl acetate and gently stirred overnight by a rotary mixer for extracting the *p-*coumaric acid. Finally, the ethyl acetate solution containing *p-*coumaric acid was subjected to evaporation in a rotary evaporator, wherein the solvent alone was removed, and the solid residue containing *p-*coumaric acid was left behind. This residue was dissolved in 0.5 M NaOH to prepare a 100 mM stock solution (pH 8.7) for production studies (Fig. S1).

### 2.7. Growth on depolymerized lignin and bioconversion of catechol to ccMA

The *p-*coumaric acid stock extracted from depolymerized lignin was diluted in mineral salt medium to achieve a final *p-*coumaric acid concentration of 15 mM, pH 7.5. Shake flask studies with *A. baylyi GJS*_*catA* ADP1 were done in duplicates in 100 ml flasks (10 ml cultures). Samples were collected periodically to monitor cell density (OD_600_) and for HPLC analysis to quantify catechol and ccMA. To check the optical density, cell pellets were subjected to two successive washing steps with 0.9% (w/v) sodium chloride solution to eliminate potential spectrophotometric interference from catechol or lignin in the growth medium.

### 2.8. Co-culture based bioconversion of depolymerized lignin to ccMA

*A. baylyi* GJS2_*catA* were grown in 100 ml shake flasks (10 ml cultures), with 5 mM glucose and 3 mM *p-*coumaric acid, extracted from depolymerized lignin. 10 OD_600_ of *E.coli*_*aroY* was added to the media as soon as all the *p-*coumaric acid was consumed and PCA accumulated. Samples were collected at frequent intervals to monitor biomass and substrate, and product concentrations.

## 3. RESULTS

### 3.1. Bioconversion of Catechol to *cis, cis* - Muconic acid

This study promises to investigate the bioconversion of depolymerized lignin, predominantly containing p-coumaric acid, to ccMA by *A. baylyi* ADP1. Before carrying out the detailed investigations on this conversion, we investigated the conversion of catechol to ccMA. In the b-ketoadipate pathway of *A. baylyi* ADP1, ccMA is produced as an intermediate metabolite in the catechol branch (Fig. 2). This necessitates optimal bioconversion of catechol to ccMA. Before optimizing the catechol to ccMA bioconversion, toxicity assays were carried out for catechol and ccMA to establish the maximum non-inhibitory substrate concentration and the potential for product inhibition. On this basis, suitable initial concentrations and feeding strategies were selected to avoid catechol toxicity.

**Figure 2.**
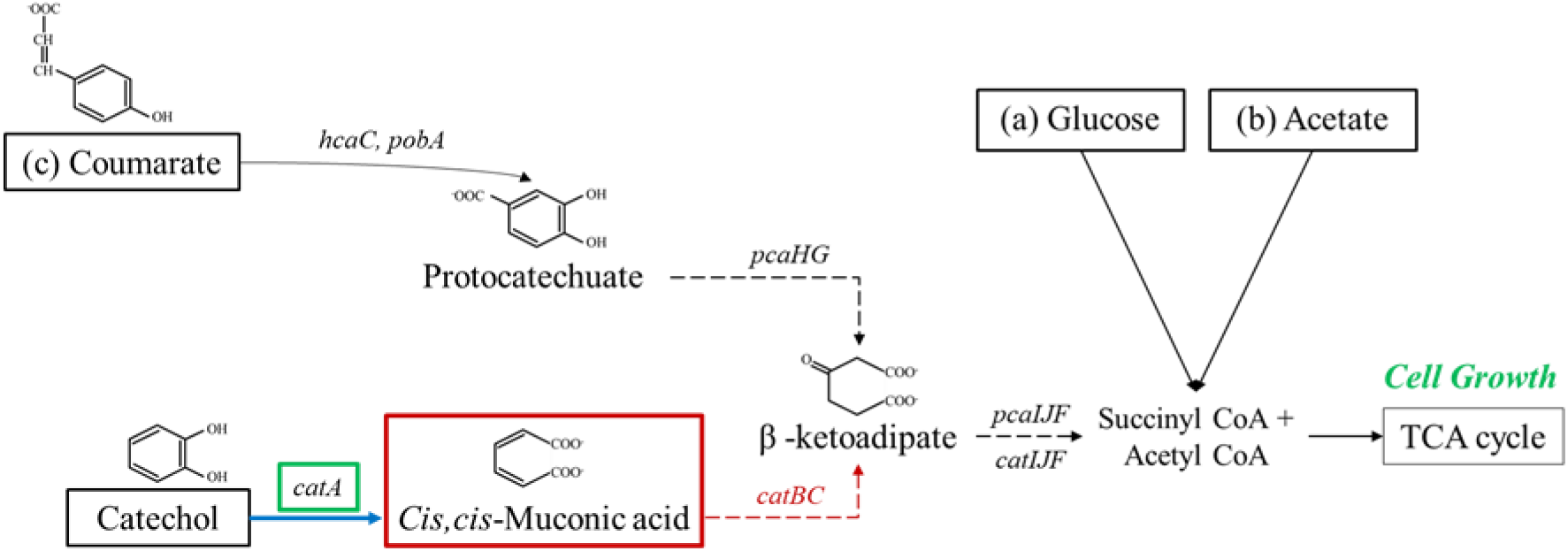
Engineered pathway in *A. baylyi* ADP1 for accumulation of ccMA from catechol in the presence of three different primary carbon sources - (a) Glucose, (b) Acetate, (c) *p*-Coumaric acid. The *catB* and *catC* genes encode muconate cycloisomerase (EC 5.5.1.1) and muconolactone Δ-isomerase (EC 5.3.3.4), respectivelyThe *catA* gene (ACIAD1442) encodes catechol 1,2-dioxygenase (EC 1.13.11.1), which is responsible for conversion of catechol to ccMA.

The results demonstrated that catechol concentrations exceeding 10 mM was inhibitory to the growth, establishing a clear toxicity threshold for this aromatic substrate (Fig. S2). The concentration-dependent inhibition is likely due to the compound’s dihydroxyl structure and associated redox properties that render it highly reactive and potentially cytotoxic (Aravind et al., 2021) This suggests that catechol accumulation during a fermentation process, beyond the cellular detoxification capacity, would result in growth inhibition. In contrast, ccMA did not demonstrate inhibitory effects on bacterial growth, even at the highest tested concentration of 60 mM (Fig. S3). This tolerance can be attributed to ccMA’s role as a natural intermediate in the β-ketoadipate pathway, where bacterial cells possess efficient enzymatic mechanisms for its metabolism (Draths & Frost, 1994). This toxicity assessment establishes that successful catechol-to-ccMA bioconversion will be primarily constrained by toxicity of catechol as substrate rather than product accumulation. The catechol-toxicity necessitates controlled feeding strategies to prevent accumulation beyond the 10 mM threshold. Conversely, the absence of product inhibition by ccMA suggests that high-titer production can be achieved without metabolic constraints or feedback inhibition effects.

We investigated the conversion of catechol to ccMA by engineering the *A. baylyi* Δ*IS* ADP1 strain. This required knockout of the *catB* and *catC* genes (Fig. 2), which prevents the downstream catabolism of ccMA through the β-ketoadipate pathway. To enhance the bioconversion efficiency of catechol to ccMA, we overexpressed the homologous *catA* gene . The *catA* gene was overexpressed by the plasmid pBWB162 containing *lacI*-Trc inducible promoter. The overexpression of the *catA* gene in *A. baylyi* GJS_*catA* strain exhibited a threefold increase in bioconversion (Fig. S4) and the accumulation of ccMA as a terminal metabolite.

We explored for bioconversion of catechol to ccMA by *A. baylyi* GJS_*catA* in dual-substrate experiments which decoupled growth and product formation. Three different substrates, glucose, acetate, and *p*-coumaric acid, were independently explored for growth, while catechol was added as a secondary substrate for ccMA production (Fig. 2).

Shake-flask cultivations were conducted in 250 mL Erlenmeyer flasks containing 25 mL of mineral salt medium at 30°C and 250 rpm. Three different primary carbon sources were evaluated: glucose, acetate, and *p-*coumaric acid. For the glucose-substrate experiments, 20 mM glucose was the initial concentration at inoculation, followed by an additional 20 mM glucose feed at 9^th^ hour, to obtain an OD_600_ of 9-10 (Fig. S10 (a)). For acetate-substrate experiments, cultures were grown with an initial concentration of 50 mM sodium acetate, with further additions of 50 mM, 40 mM, and 40 mM at 9^th^, 15^th^ and 21^st^ hour, respectively; this feeding schedule supported growth to an OD_600_ of 6-7 (Fig. S10 (b)). In the third study, *p-*coumaric acid was employed as the primary feed, with an initial supplementation of 15 mM followed by sequential additions of 15 mM and 10 mM at 12^th^ and 15^th^ hour, to sustain growth to an OD_600_ of 9-10 (Fig. S10 (c)).

Upon reaching maximum biomass, 10 mM catechol was added every 3 h to drive ccMA synthesis. The primary carbon source was used to support cellular growth in the presence of catechol. Comparative analysis of growth kinetics and substrate consumption profiles revealed that *p-*coumaric acid as primary carbon source supported the highest ccMA titres, achieving 44.65 ± 0.11 mM. Glucose and acetate experiments however yielded lower product concentrations of 15.65 ± 0.39 mM and 27.82 ± 0.52 mM, respectively (Fig. S10). In all the experiments, catechol consumed was converted to ccMA with an approximate carbon yield of 0.9 C-mmol ccMA per C-mmol catechol. These results indicate that *p-*coumaric acid is the preferred primary carbon source for facilitating best catechol to ccMA production in the dual-substrate cultivation of *A. baylyi* GJS_*catA* strain, as compared to acetate and glucose.

To explore the scalability of this process, *A. baylyi* GJS_*catA* strain was cultivated in a 1.2 L stirred-tank bioreactor. Since step-feeding of catechol in the shake-flask cultures resulted in catechol accumulation, we investigated the impact of continuous substrate feeding on ccMA production in the bioreactor. Catechol was added at a constant rate of 2.5 mM h⁻¹ via peristaltic pump, minimizing transient substrate accumulation and associated toxicity. This feeding strategy yielded a mean ccMA titre of 50.04 ± 6.31 mM across duplicate experiments, with one reactor achieving a maximum of 56.4 mM (Fig. 3). These values exceed those obtained under step*-*wise feeding conditions in shake-flask cultures, demonstrating that gradual substrate supply enhances product formation by avoiding the initial inhibitory effects of high catechol concentration during the stepwise addition.

**Figure 3.**
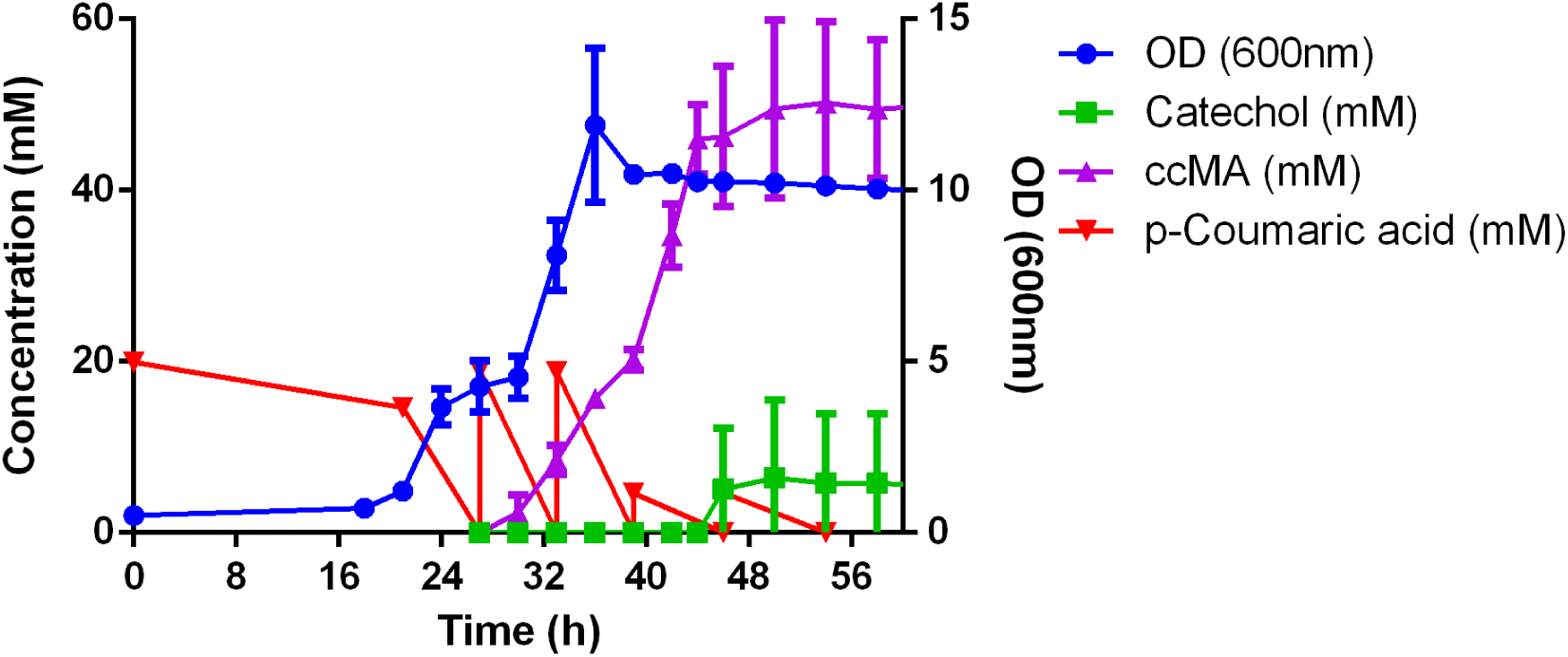
Growth of *A. baylyi A. baylyi* GJS_*catA* ADP1 in *p-*coumaric acid (as primary carbon source for growth) and catechol (as secondary carbon source for ccMA production) by in a 1.2 L reactor, with continuous feeding of catechol at 2.5 mM per hour rate.

**Figure 4.**
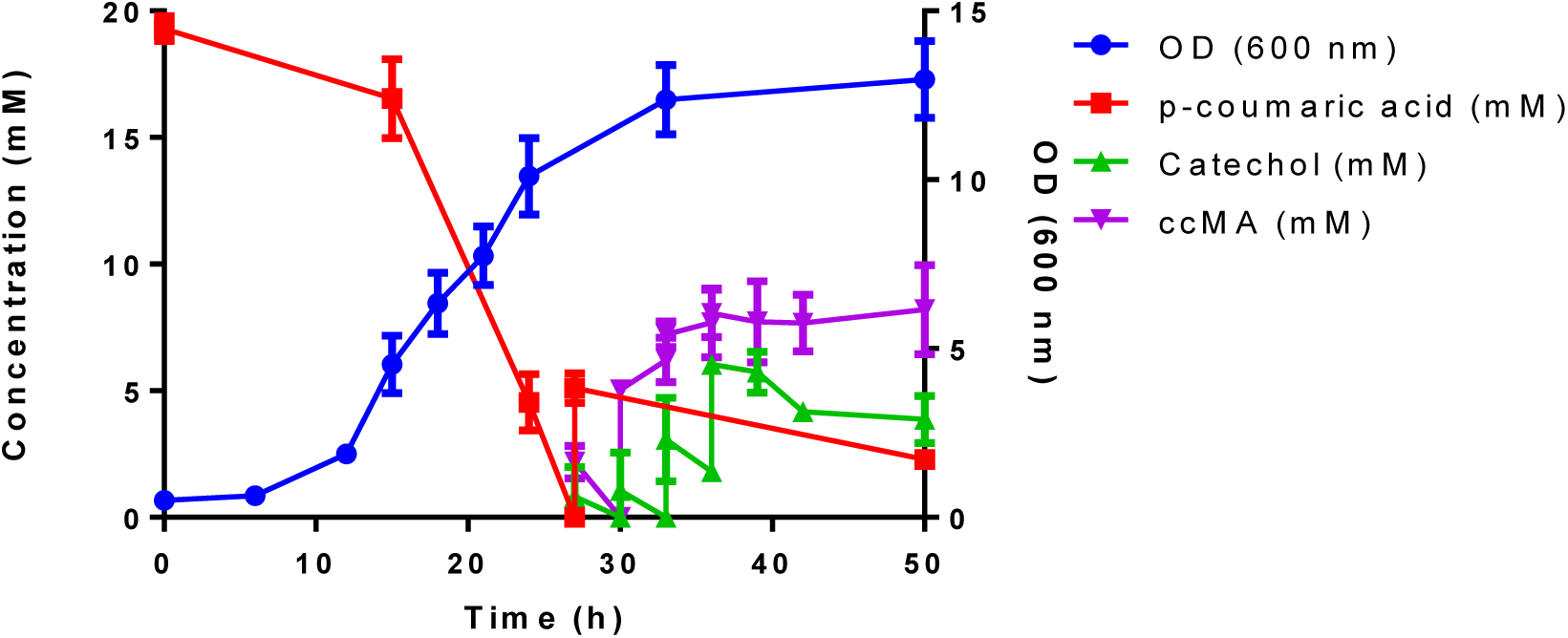
Growth in depolymerized lignin and catechol to ccMA bioconversion by *A. baylyi* GJS_*catA* ADP1 in a 10 ml shake-flask cultures.

### 3.2. Growth studies on depolymerized lignin and bioconversion of catechol to ccMA

In the next set of studies, we evaluated the feasibility of using depolymerized lignin as feedstock for ccMA production. Depolymerized lignin fractions were derived from alkali-treated lignin-rich residue of pre-treated, enzymatically hydrolysed corncob biomass (Fig. S1). These depolymerized fractions predominantly contained p-coumaric acid which was further purified by extraction with ethyl acetate and 20 mM employed as the primary carbon source for cultivation of *A. baylyi* GJS_*catA* strain. Catechol was added separately for bioconversion to ccMA. After 24 hours, when the cells reached > 10 OD_600_, 4 mM catechol was added to the media, at an interval of 3 hours from 27^th^ hour (12 mM of catechol in total). The final ccMA titer was 8.42 ± 1.02 mM, with a carbon yield of approximately 0.9 C-mmol ccMA per C-mmol catechol (accounting for a residual catechol of 3.87 ± 0.65 mM) (Fig. 5). These results demonstrate successful catechol bioconversion to ccMA in the presence of depolymerized lignin as the sole primary carbon source, confirming the potential of *A. baylyi* ADP1 to grow on depolymerized lignin.

**Figure 5.**
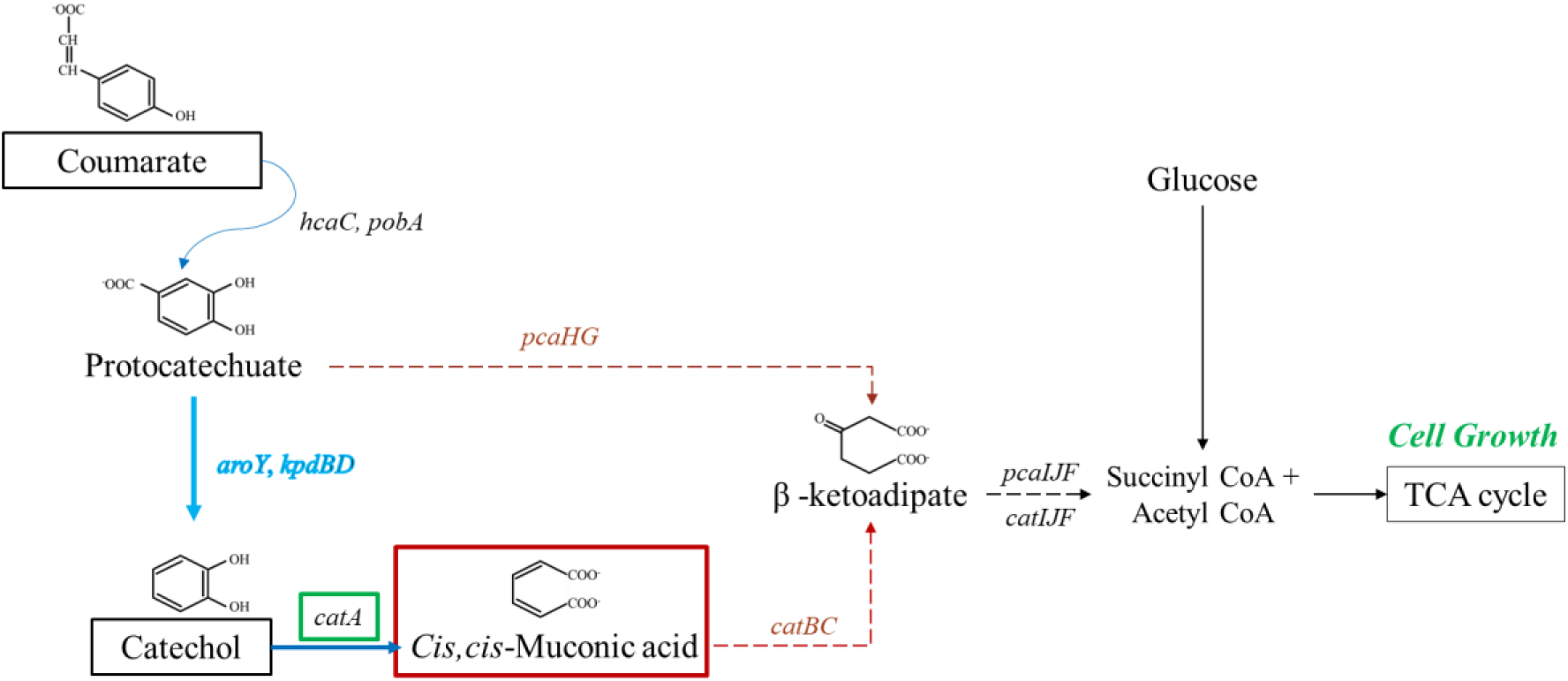
Schematic pathway for bioconversion of *p-*coumaric acid to ccMA with glucose as growth substrate in *A. baylyi* ADP1. The *pcaHG* genes encode for PCA 3,4-dioxygenase beta and alpha (1.13.11.3).

### 3.3. Bioconversion of *p-*coumaric acid to ccMA

Having validated the bioconversion of catechol to ccMA, the ability of the strain to convert *p*-coumaric acid to ccMA was investigated. This requires conversion of the key intermediate PCA to catechol by PCA decarboxylase, which is missing in *A. baylyi* and needs to be heterologously expressed (Fig. 5). To enable directed conversion of PCA to catechol, other genetic modifications were required to accumulate PCA and prevent further catabolism in the b-ketoadipate pathway.

Targeted genetic modifications were performed on the *A. baylyi* GJS strain to enable PCA accumulation from *p-*coumaric acid. The genes *pcaHG* genes were deleted, thus disabling the further metabolism of PCA in the b-ketoadipate pathway (Fig. 5). For further pathway optimization, the *acr1* gene was knocked out. This gene codes for fatty acyl-CoA reductase, which helps in wax ester production (1.2.1.42). The *acr1* gene is responsible for wax ester biosynthesis in *A.baylyi* ADP1, and diverts acetyl-CoA and NADPH towards storage lipid accumulation. Disruption of this pathway is expected to enhance carbon flux toward central metabolism and downstream product formation such as ccMA (Arvay et al., 2021). The resultant strain was named *A. baylyi* GJS2 (Table S2).

The codon-optimized *aroY* gene that codes for PCA decarboxylase, along with *kpdBD* genes, was incorporated into the *A. baylyi* GJS2 strain using the plasmid pBWB162. These genes were taken from *Klebsiella pneumoniae,* where the *kpdB* gene codes for FMN co-reductase and *kpdD* stabilizes the full complex. In a previous study, these genes were incorporated in *Pseudomonas putida* KT2440, where it was seen that incorporation of the *kpdB* gene along with the *aroY* gene increased the decarboxylation efficiency (Sonoki et al., 2014). Hence, the strain *A. baylyi* GJS2_*aroYkpdBD* was constructed and tested for bioconversion of PCA to catechol. However, in a whole-cell biotransformation assay, it was observed that the *A. baylyi* GJS2_*aroYkpdBD* strain did not show any bioconversion of PCA to catechol (Fig. S9). qPCR analysis was conducted to assess the transcription of the codon-optimized *aroY* gene in *A. baylyi* ADP1. Comparative expression profiling between strains *A. baylyi* GJS2 and *A. baylyi* GJS2_*aroYkpdBD* strains confirmed successful transcription of the codon-optimized *aroY* construct. Despite this, functional protein activity was absent (Fig. S5). Recent findings by C. Liu et al. (2025) indicate that the inability of engineered *A. baylyi* strains to perform the targeted bioconversion may result from the presence of additional regulatory genes such as *benR* and *catM*.

*E.coli* strain was also engineered to check the activity of the heterologous *aroY* gene to perform the bioconversion from PCA to catechol. *E.coli*_*aroY* strain, containing the wild-type *aroY* gene, was easily able to convert PCA to catechol.

This prompted the design of a co-culture strategy wherein the PCA to catechol conversion step will be taken care of by the engineered *E.coli* strain. Since the wild-type *aroY* gene showed better bioconversion as compared to the codon-optimized genes in *E.coli*, *E.coli*_*aroY* strain was selected for co-culture with *A. baylyi* GJS2_*catA* strain, where the catechol decarboxylase gene has also been overexpressed. There are 3 steps for the bioconversion of *p-*coumaric acid to ccMA in this process (Fig. 6): *p-*coumaric acid to PCA bioconversion by *A. baylyi* GJS2_*catA* strain; bioconversion of accumulated PCA to catechol by *E.coli*_*aroY*; and finally catechol to ccMA conversion by *A. baylyi* GJS2_*catA* strain.

**Figure 6.**
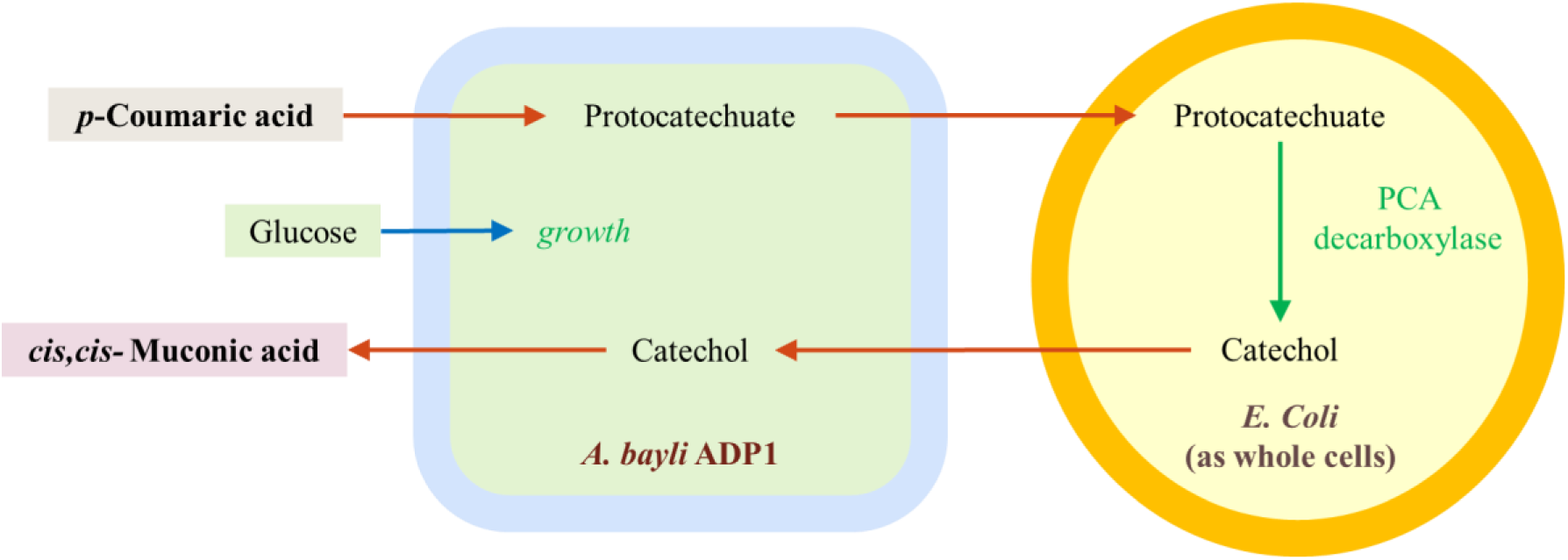
Schematic pathway for bioconversion of *p-*coumaric acid to ccMA with glucose as growth substrate with co-culture.

This process would require the accumulation of PCA from *p*-coumaric acid in the first step. It was observed that *A. baylyi* GJS2_*catA* strain was able to utilize *p-*coumaric acid as the sole carbon source and grow on it (Fig. S6). There was no PCA accumulation from *p-*coumaric acid. This meant that the *pcaHG* knockout does not prevent further metabolism of PCA and needs additional gene deletions for the accumulation of PCA (Liu et al., 2025). On the other hand, when glucose was added as a primary carbon source for growth and *p-*coumaric acid as a secondary substrate for ccMA production, it was observed that there was transient PCA accumulation, which was later completely depleted (Fig. S7). To improve the PCA accumulation, step*-*feed of incremental amounts of glucose and *p-*coumaric acid would be desirable.

A whole-cell biotransformation study of *p-*coumaric acid with both *A. baylyi* GJS2_*catA* and *E.coli*_*aroY* strains showed gradual conversion of the substrate to its intermediate metabolites, i.e., 4-hydroxybenzoic acid, PCA, and finally to ccMA (Fig. S8). The rate of biotransformation was very slow, yielding only 15% successful bioconversion to ccMA in 10 hours. Although *p-*coumaric acid was completely consumed, intermediate metabolites like 4-hydroxybenzoic acid were accumulated, indicating that a faster bioconversion of *p-*coumaric acid to PCA is necessary for efficient ccMA production. Hence, accumulation of PCA before the addition of *E.coli*_*aroY* was explored in the next set of studies.

To accumulate PCA before the addition of *E.coli*_*aroY* strain, *A. baylyi* GJS2_*catA* strain was grown in media with a step-feed of 5 mM glucose for growth and 3 mM *p-*coumaric acid for accumulating protocatechuate (Fig. 7). It was observed that *A. baylyi* GJS2_*catA* strain was able to accumulate around 20.65 ± 0.34 mM of PCA from ∼22.1 mM of *p-*coumaric acid in 22 hours, indicating high bioconversion. Upon addition of 10 OD_600_ of *E.coli*_*aroY* strain, it yielded around 18.55 ± 0.54 mM of ccMA in 14 hours (Fig. 9). This indicates that the co-culture strategy can successfully convert *p-*coumaric acid to ccMA when the feeding of the carbon sources is optimized. The C/C yield of ccMA from *p-*coumaric acid is 0.56, corresponding to 83.6% of the theoretical maximum (0.67).

**Figure 7.**
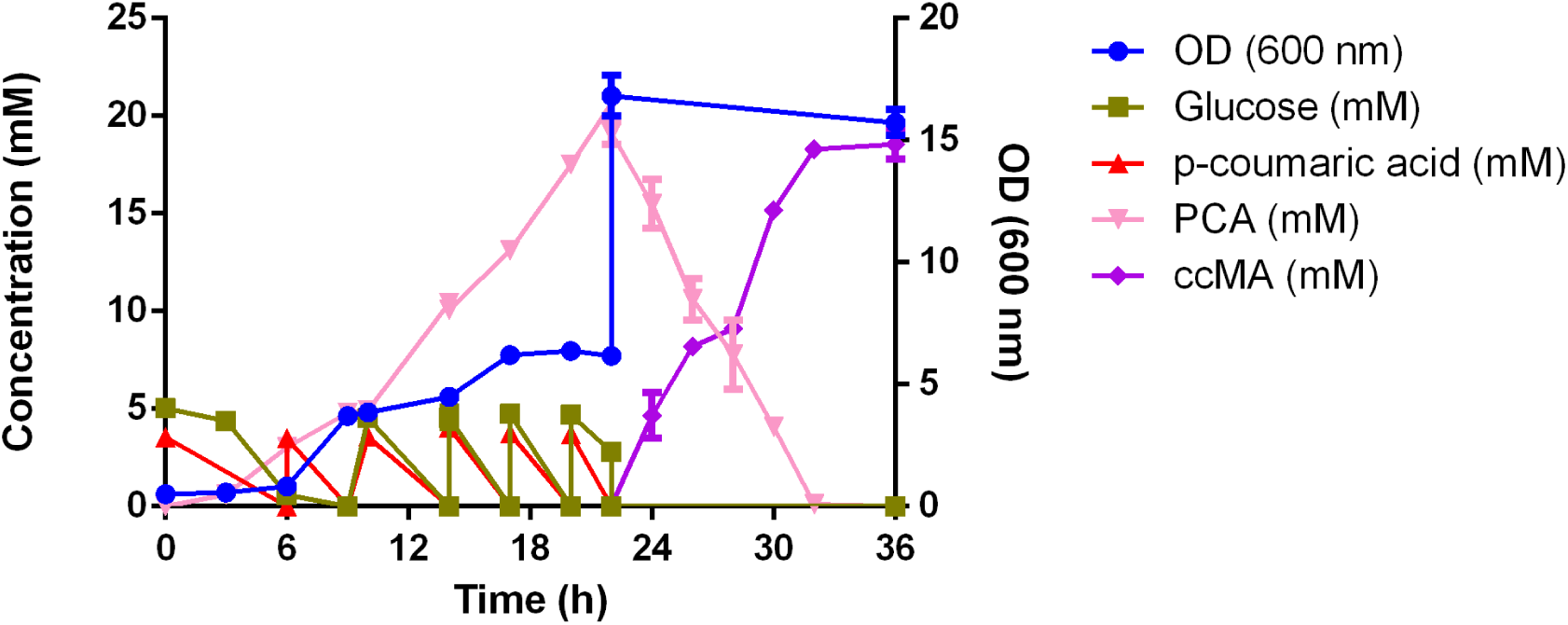
Shake flask study of co-culture in *p-*coumaric acid and glucose feed

In the present study, a final titer of 18.55 ± 0.54 mM ccMA (equivalent to 2.63 g/L) was achieved. This substantial enhancement in product yield was realized through the implementation of a co-culture system, specifically by pairing engineered *E. coli* with *A. baylyi* ADP1 and deleting the wax ester pathway gene (*acr1*).

### 3.4. Bioconversion of depolymerized lignin to ccMA

In the final set of studies, the bioconversion of depolymerized lignin to ccMA was explored. *A. baylyi* GJS2_*catA* strain was grown in media supplemented at 0^th^ and 12^th^ hour with 4 mM *p-*coumaric acid extracted from depolymerized lignin. In addition, 5 mM glucose was added at 0^th^ and 9^th^ hour (Fig. 8). It took approximately 9-12 hours for the conversion of ∼5 mM *p-*coumaric acid to PCA, compared to 4-6 hours taken for the conversion of synthetic *p-*coumaric acid. From 8.35 mM of *p-*coumaric acid, 7 mM PCA was accumulated in 21 hours, showing 85% conversion of *p-*coumaric acid to PCA. However, upon addition of 10 OD_600_ of *E.coli*_*aroY* strain at 21^st^ hour, only 1.75 ± 0.02 mM ccMA was produced in 26 hours, showing only 24% conversion (Fig. 10). The rest of the PCA was metabolized by *A. baylyi* GJS2_*catA* strain. The inhibitory effects of the depolymerized lignin fraction on *E. coli* limited PCA to catechol production and subsequent ccMA synthesis. Conversely, the resilience of *A. baylyi* ADP1 in lignin-containing media allowed it to metabolize residual PCA effectively. This proof-of-concept study demonstrates the potential for co-culture-based valorization of depolymerized lignin but indicates the need for optimization to improve titers and yields of ccMA.

**Figure 8.**
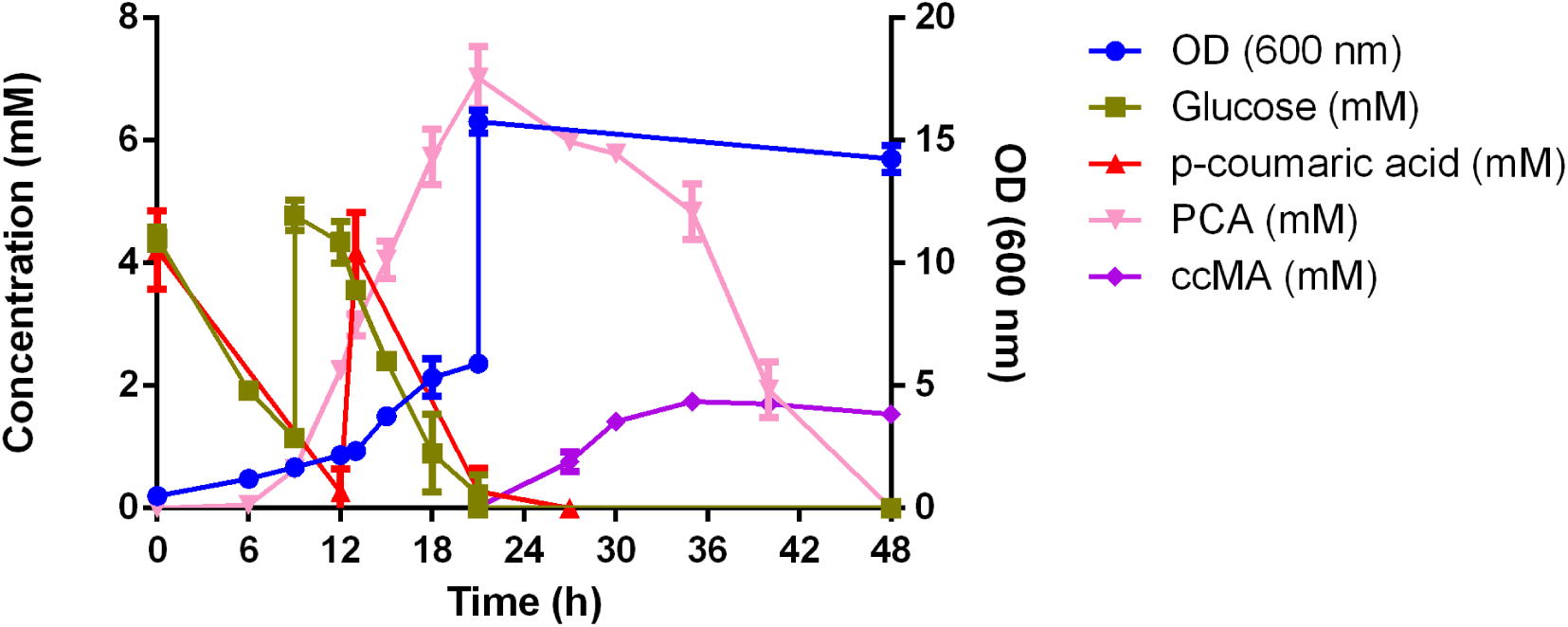
Growth in glucose and depolymerized lignin to ccMA bioconversion by co-culture in 2-steps in a 10 ml shake-flask cultures. First step - 0-22 hours (PCA accumulation from Cm in depolymerized lignin), Second step - 22-48 hours (PCA-CA-ccMA)

## 4. DISCUSSION AND CONCLUSION

The present study establishes an engineered *A. baylyi*-*E. coli* co-culture as an effective framework for the valorisation of *p*-coumaric acid, a prominent lignin-derived monomer, to ccMA. Initial investigation focused on bioconversion of catechol to ccMA, in a dual-substrate setup. A variety of genetic engineering strategies have been developed to enable and improve the bioconversion of catechol (as an initial carbon source) to ccMA. The most prominent of them is the dual substrate strategy, wherein catechol is used specifically as the precursor for ccMA production, while a second substrate is supplied solely for cellular growth. Table 1 highlights many such studies, and the available evidence demonstrates a C/C yield as high as 0.9-0.95, as the entire catechol flux is directed through a single-step conversion to ccMA. *Corynebacterium glutamicum* has been the most efficient microbial host for the dual substrate strategy, producing the highest titer of ccMA of 85 g/L from catechol in recent studies (Becker et al., 2018).

Deletion of *catB* and *catC* to truncate the β-ketoadipate pathway, combined with *catA* overexpression in the *A. baylyi* GJS_*catA* strain substantially increased the flux from catechol to ccMA by three fold (Fig. S4). Overexpression of catechol 1,2-dioxygenase provides elevated levels of the key enzyme responsible for catechol ring cleavage, thereby increasing metabolic flux through the desired pathway while reducing substrate accumulation. This enhancement in biotransformation kinetics demonstrates the effectiveness of the combined deletion and overexpression (push-pull) strategy for optimizing ccMA production. This has been observed previously in *Pseudomonas* and *Acinetobacter* where catechol cleavage has been identified as rate-limiting for ccMA accumulation from diverse aromatic substrates (Draths & Frost, 1994). Based on these comparative results, the *A. baylyi* GJS*_catA* strain was selected for subsequent growth and production studies.

The comparison of glucose, acetate, and *p*-coumaric acid as primary substrates for the *A. baylyi* GJS*_catA* strain in the dual-substrate experiments showed that *p*-coumaric acid supported the highest biomass formation and ccMA titres (44.65 mM) relative to acetate (27.82 mM) and glucose (15.65 mM) from catechol as the secondary substrate. The strain showed similar catechol-to-ccMA yields (∼0.9 C/C) across all conditions. This finding suggests that *p*-coumaric acid, which is naturally channelled through the β-ketoadipate pathway in *A. baylyi* ADP1, provides a more favourable energetic and regulatory environment for mixed-substrate metabolism than purely glycolytic or acetyl-CoA-based feed. This is likely due to better coordination between aromatic catabolism and central carbon metabolism. When the process was transferred from stepwise catechol addition in flasks to continuous feeding in a stirred-tank reactor, ccMA titres increased further to ∼50 mM on average (maximum 56.4 mM), confirming that gradual catechol supply alleviates toxicity and improves overall conversion.

Notably, both reactor studies exhibited an extended lag phase of approximately 21 h, a phenomenon absent in previous shake-flask experiments. Such prolonged adaptation has been attributed to the physiological adjustments required under controlled bioreactor conditions, including variable oxygen transfer rates, increased foaming despite antifoam addition, and altered osmotic pressures relative to shake flasks (Brauneck et al., 2025; Rolfe et al., 2012). This study confirms that continuous catechol feeding is superior to step*-*addition for optimizing ccMA titres at reactor scale. It also highlights the need to address reactor-specific challenges such as lag-phase mitigation, foaming control, and oxygen-mass transfer optimization, to further improve process robustness and productivity.

When depolymerized corncob lignin was introduced as the source of *p*-coumaric acid, *A. baylyi* GJS*_catA* strain retained its capacity to grow and carry out catechol-to-ccMA conversion, achieving a similar carbon yield on catechol (∼0.9 C/C) but a lower ccMA titre (8.42 mM) and pronounced catechol accumulation. The extended lag phase observed initially is consistent with known physiological stress responses and metabolic adaptations required for growth on complex lignin-derived substrates, as documented in controlled culture studies of *A. baylyi* ADP1 (Brauneck et al., 2025; Rolfe et al., 2012). This adaptation period must be accounted for in designing and scaling lignin-based bioprocesses to optimize productivity and operational efficiency. Nonetheless, the data provide a proof-of-concept that *A. baylyi* GJS*_catA* strain can utilise authentic lignin-derived feeds and that the catechol to ccMA bioconversion remains functional under these harsher conditions. Future work should focus on improving oxygen transfer, refining lignin pretreatment or extract clean-up to reduce inhibitory species, and exploring adaptive evolution with depolymerized lignin to shorten lag phases and increase catechol utilisation.

The second part of the study addresses the more challenging goal of converting *p*-coumaric acid to ccMA via the PCA branch. Sequential deletions of *pcaHG* and *acr1* in *A. baylyi* GJS strain were designed to enforce PCA accumulation from *p-*coumaric acid and to eliminate the competing wax ester pathway. This strain was named *A. baylyi* GJS2 strain. Integration of heterologous *aroYkpdBD* cassette from *K. pneumoniae*, in *A. baylyi* GJS2 strain failed to convert PCA to catechol. However, engineered *E. coli* strain with the *aroY* gene readily converted PCA to catechol under the same conditions. These contrasting outcomes underscore that the protein activity is strongly host-dependent and that *E. coli_aroY* strain is currently a more suitable chassis for this specific decarboxylation step than *A. baylyi*. They also suggest that further regulatory or chaperone engineering would be required to realise an effective PCA decarboxylase in *A. baylyi*, potentially including additional deletions (e.g. *benR*, *catM*) as suggested by recent literature (Liu et al., 2025).

Given these limitations, this study investigated a co-culture strategy that distributed pathway tasks between *A. baylyi* and *E. coli*. In the designed scheme, *A. baylyi* GJS2_*catA* strain converts *p*-coumaric acid to PCA, *E. coli*_*aroY* strain decarboxylates PCA to catechol, and again *A. baylyi* GJS2_*catA* strain completes the route by converting catechol to ccMA via overexpressed *catA*. However, there were two bottlenecks in this process. First, PCA was still being metabolized in *A. baylyi* GJS2_*catA* strain despite knocking out downstream genes for PCA accumulation. This behaviour is consistent with the presence of alternative ring-cleavage routes or regulatory mechanisms that maintain flux from PCA into central metabolism (Liu et al., 2025). Thus, the *A. baylyi* GJS2_*catA* strain requires additional metabolic engineering to prevent further PCA catabolism. Secondly, the presence of *p*-coumaric acid in the media hinders the PCA to catechol conversion by the *E. coli*_*aroY* strain, as seen in the whole-cell biotransformation study (Fig. S8).

Hence, a growth study for maximal accumulation of PCA from *p*-coumaric acid prior to addition of *E. coli*_*aroY* strain as whole cells was investigated as a better alternative. The addition of incremental amounts of both glucose and *p*-coumaric acid proved beneficial for the successful accumulation of PCA before the addition of the *E. Coli* strain. When > 5 mM of *p*-coumaric acid was added to the media, the conversion to PCA was either slow or inefficient (Fig. S7). When > 5 mM of glucose was added, the *A. baylyi* GJS2_*catA* strain took longer to metabolize it, which would lead to presence of glucose in the media when the *E.coli_aroY* strain is added. Hence, a step-feed of 3 mM of *p*-coumaric acid and 5 mM of glucose was optimized and resulted in better ccMA titers. The co-culture shake-flask experiments addressed the kinetic mismatches by decoupling PCA accumulation from its downstream conversion. Under a carefully optimised step-feed of glucose (for growth) and *p*-coumaric acid (for PCA formation), *A. baylyi* GJS2_catA strain accumulated ∼20.65 mM PCA from ∼22.1 mM p-coumaric acid within 22 h. Subsequent addition of *E. coli_aroY* strain led to the production of 18.55 mM ccMA in 14 h, corresponding to a 83.6% C/C yield of the theoretical maximum. This is a substantial improvement over earlier *A. baylyi*-based ccMA processes, and demonstrates that division of labour between an aromatic degrader and a dedicated PCA decarboxylation partner can be highly effective. Notably, this performance was achieved without further gene deletions in *A. baylyi*, suggesting that co-culture design can, at least in part, substitute for deeper regulatory rewiring when appropriate process control is applied.

The final experiments, in which *p-*coumaric acid derived from depolymerized lignin was used instead of the synthetic aromatic, revealed important constraints for lignin-based co-cultures. This study exposed the complexities of multi-organism systems in lignocellulosic valorization. While *A. baylyi* showed resilience and could effectively convert *p-*coumaric acid derived from lignin to PCA, the *E. coli* strain engineered for PCA to catechol conversion was inhibited by depolymerized lignin fraction. This limited the overall ccMA yield. This experiment highlights the challenge of microbial compatibility in co-culture systems using complex substrates and suggests that further strain engineering, like adaptive laboratory evolution, may be necessary to maximize synergies and product yield. A setup where the *E. coli* strain has been immobilized, and there is a continuous feed of PCA produced and accumulated by *A. baylyi* GJS2_catA strain, can also be explored for the same. The catechol produced by the immobilized *E.coli* system is then fed back to the *A. baylyi* culture for ccMA production.

Overall, these findings substantiate *A. baylyi* ADP1 as a promising chassis for lignin valorization through metabolic engineering and process integration. The ability to convert authentic lignocellulosic waste streams, rather than model substrates, shows practical applicability. Continued efforts to refine metabolic pathways, optimize substrate feeding and culture conditions, and improve co-culture compatibility are essential next steps to develop economically viable bioconversion platforms for the sustainable production of industrial chemicals such as ccMA.

## List of Abbreviations

ccMA: *cis, cis* – Muconic acid
Cm: *p* - Coumaric acid
Hb: 4-hydroxybenzoic acid
HPLC: High Performance Liquid Chromatography
IS: Insertion sequence
LRAs: Lignin-related aromatics (referring to synthetic aromatics)
MSM: Minimal salt medium
OD: Optical density at 600 nm
PCA: Protocatechuic acid
PCR: Polymerase chain reaction

## Consent for publication

This work has been consented by the Department of Biotechnology, IIT Madras, India, and the Institute of Applied Microbiology, RWTH Aachen University, Germany.

## Availability of data and materials

All the remaining data are mentioned in the supporting information file.

## Authors’ Contributions

GJ conceptualized the project; SM contributed to the experimental design and performed the experiments, analytics and growth studies on depolymerized lignin; TP contributed to lignin de-polymerization and *p*-coumaric acid extraction; GJ and SM wrote the manuscript; GJ was the Principal Investigator for this project and recipient of the project funding; LMB was the co-Principal Investigator for this project. All authors read and approved the manuscript.

## Acknowledgement

We thank Prof. J. E. Barrick, University of Texas, Austin, for kindly providing us with IS1236-deleted *Acinetobacter baylyi* ADP1 strain, and Dr. Vasanthkumar Sagar from Sparconn Life Sciences, Bangalore, India for providing us the corn cob.

