## Supplementary Information for "Bioconversion of *p*-coumaric acid to *cis,cis*-muconic acid using an engineered *A. baylyi* ADP1 - *E. coli* co-culture"

### Figures:

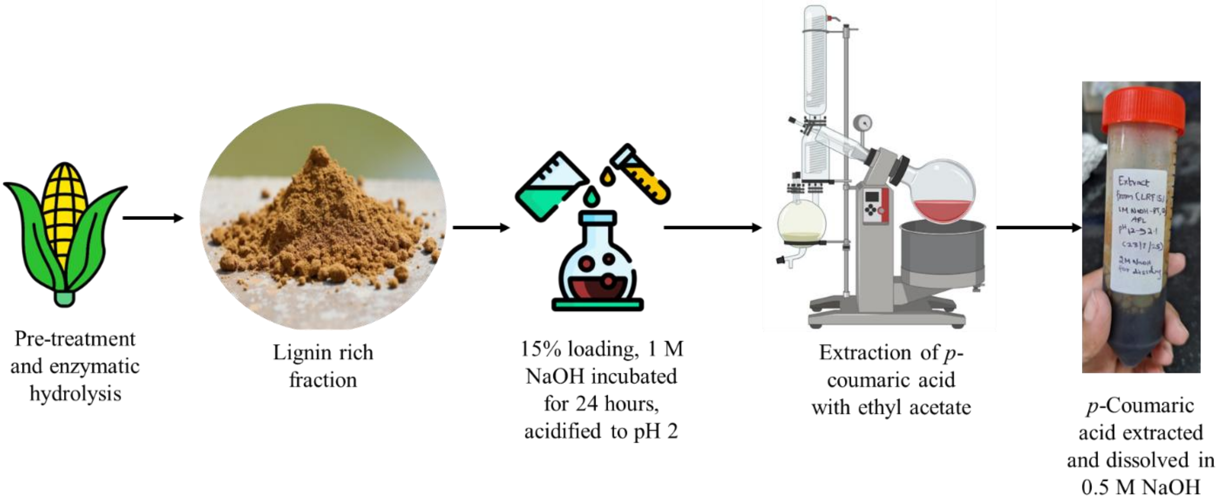


**Figure S1.** *p*-coumaric acid extracted from alkali-pretreated lignin.

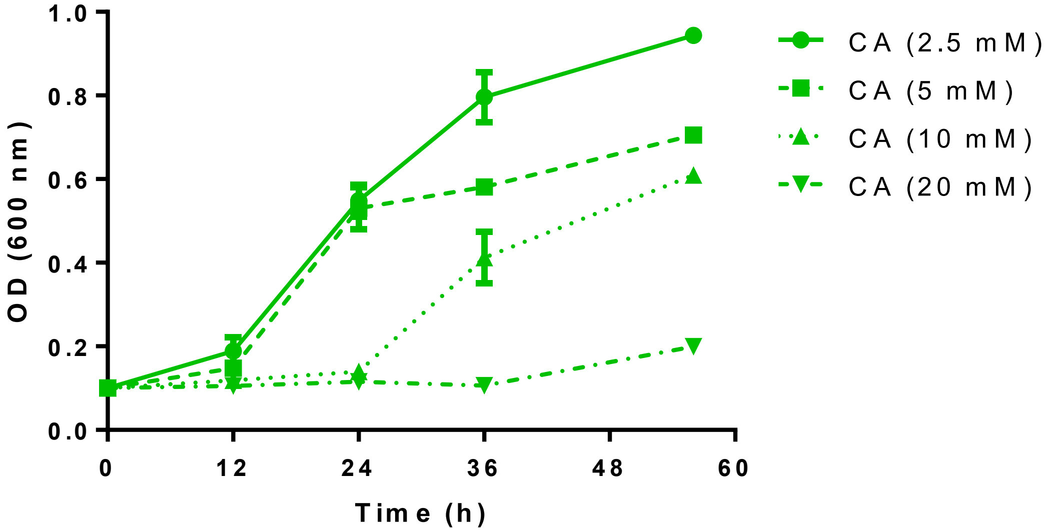


**Figure S2** Growth profiles of *A. baylyi* Δ*IS* ADP1 on different concentrations of catechol (CA).

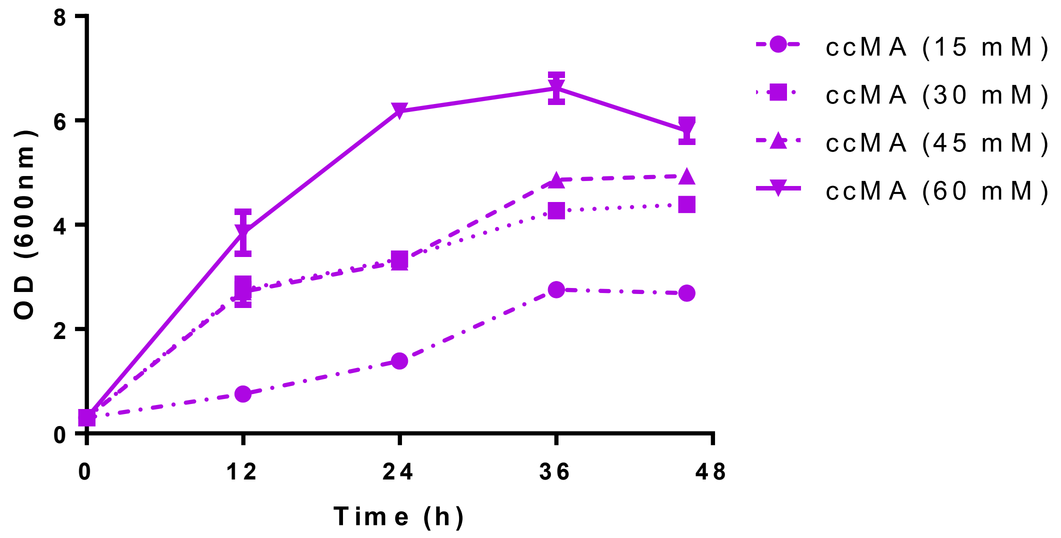


**Figure S3** Growth profiles of *A. baylyi* Δ*IS* ADP1 on different concentrations of ccMA.

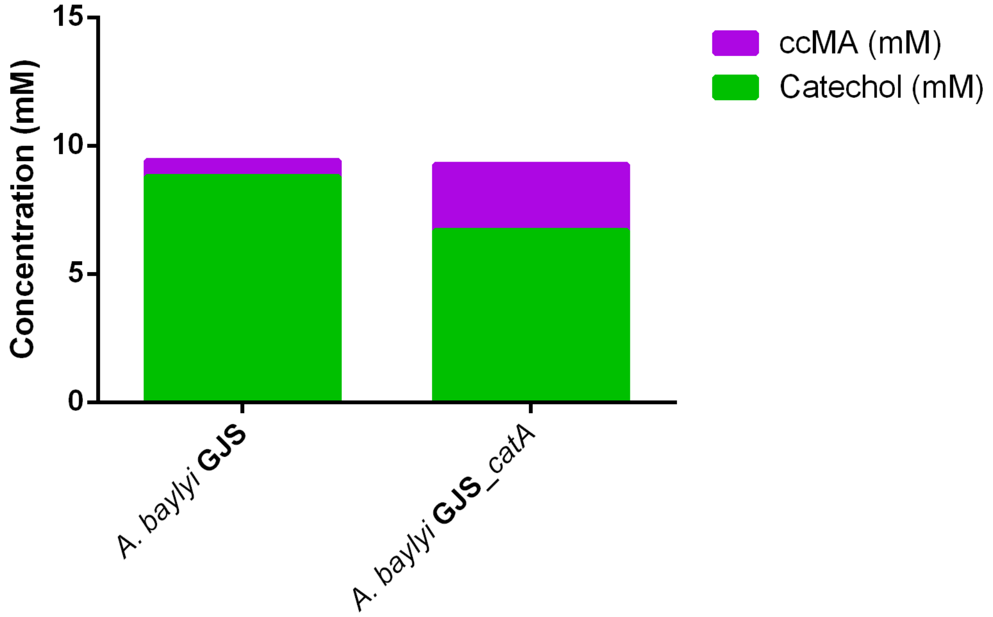


**Figure S4** Whole-cell biotransformation assay of 1 $OD_{600}$ of cells in 1 hour, for 10 mM of catechol.

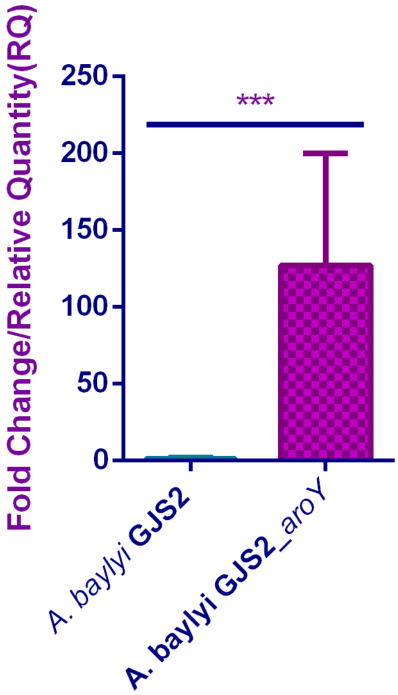


**Figure S5** Fold change of mRNA for codon-optimized *aroY* in engineered *A. baylyi* Δ*IS* ADP1 strain.

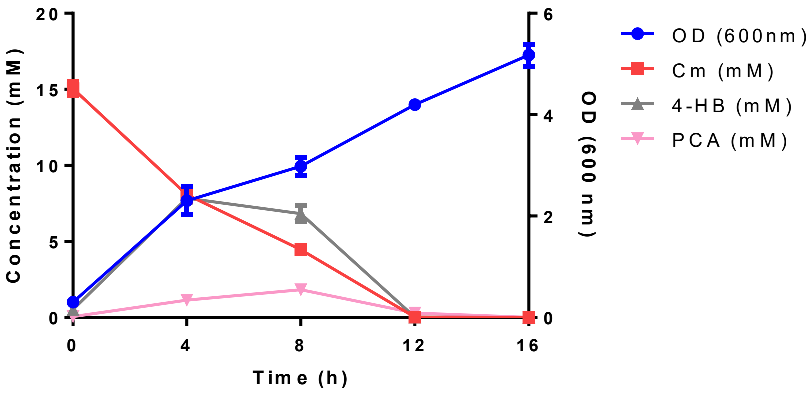


**Figure S6** Growth of *A. baylyi* GJS2 in 15 mM *p*-coumaric acid.

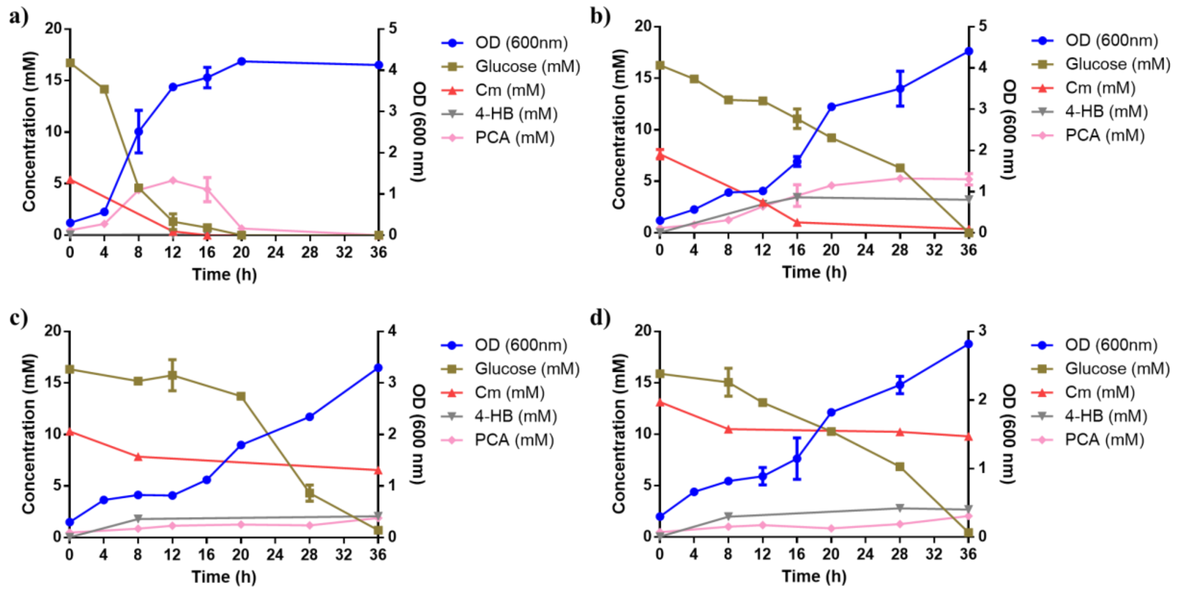


**Figure S7** Growth of *A. baylyi* GJS2 in 15 mM glucose supplemented for growth and (a) 5 mM, (b) 7.5 mM (c) 10 mM and (d) 12.5 mM *p*-coumaric acid for PCA production.

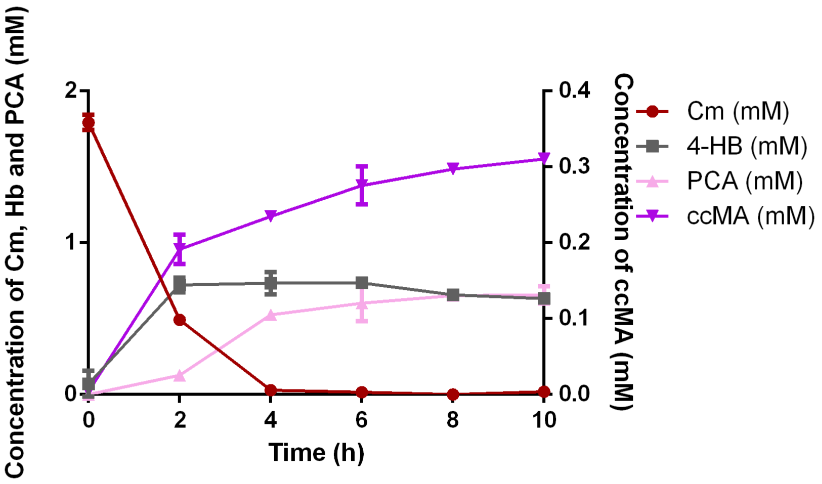


**Figure S8** Biotransformation trend of 1 mM of *p*-coumaric acid to ccMA using co-culture of the *A. baylyi* GJS2_*catA* and *E.coli_aroY* strains.

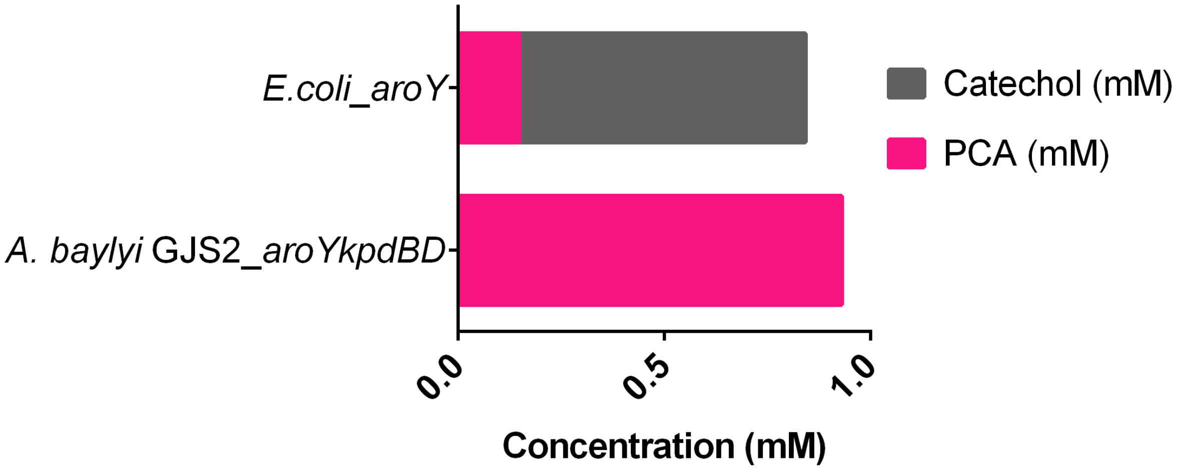


**Figure S9** Biotransformation of 10 $OD_{600}$ of the strains in 1 mM protocatechuate after 3 hours.

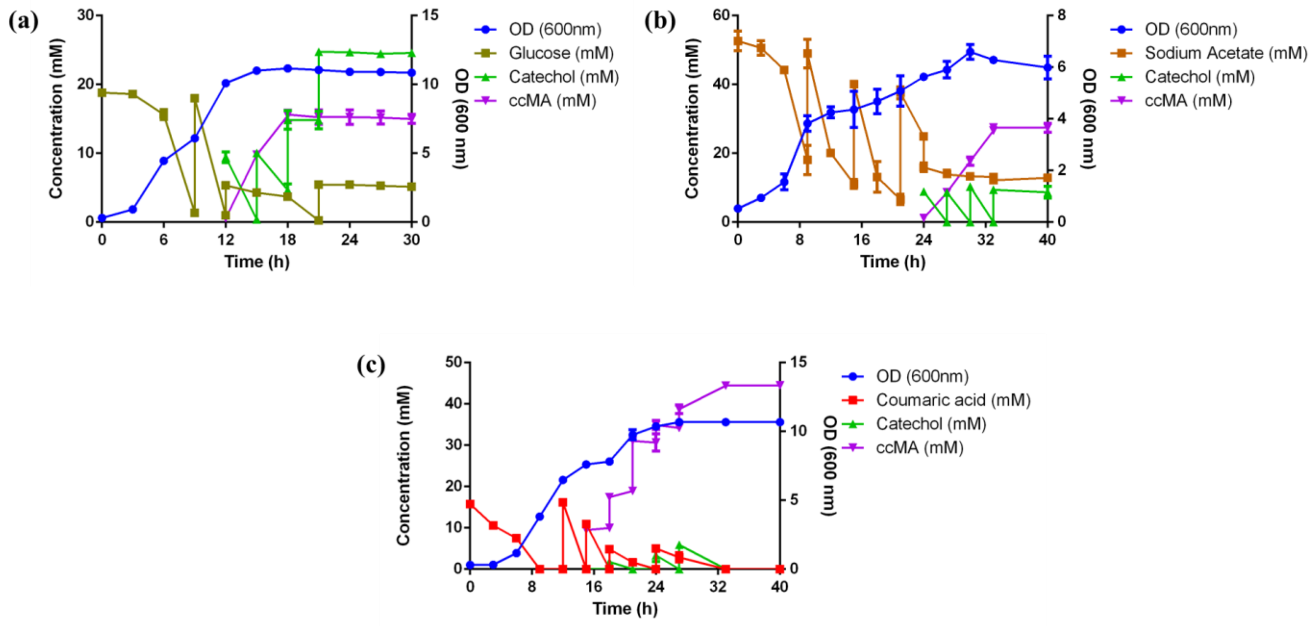


**Figure S10** Growth and substrate utilization of *A. baylyi* GJS_*catA* ADP1 in (a) Glucose and catechol, (b) Acetate and catechol, and (c) *p*-coumaric acid as primary substrate, and catechol as a secondary substrate for ccMA production, in shake flasks. Catechol was fed in a ste*p*-wise fashion.

### Tables in this study:

**Table S1** List of plasmids constructed for this study.

| Sl. No. | Plasmid name | Description |
|---|---|---|
| 1 | pBWB162_*catA* | Homologous overexpression of the *catA* gene from the *A. baylyi* ADP1 genome in the pBWB162 plasmid |
| 2 | pBWB162_*aroY* | pBWB162 with wild-type *aroY* gene (NCBI Accession No. AB479384), which has been taken from *K. pneumoniae* |
| 3 | pBWB162_*aroY*$^c$ | pBWB162 with *aroY* gene codon-optimized for *A. baylyi* ADP1 |
| 4 | pBWB162_*aroYkpdBD* | pBWB162 with codon-optimized *aroY* and *kpdBD* genes (NCBI Accession No. – AB920346), where *kpdBD* genes have been taken from *K. pneumoniae* |

**Table S2** List of strains generated in this study.

| Sl. No. | Strain Name | Description |
|---|---|---|
| 1 | *A. baylyi* GJS ADP1 | *A. baylyi ΔISΔcatBC* ADP1, where the *catBC* genes have been knocked out, unable to grow on catechol. |
| 2 | *A. baylyi* GJS_*catA* ADP1 | Overexpression of the *catA* gene in pBWB162 for efficient bioconversion of catechol to ccMA. |
| 3 | *A. baylyi* GJS2 ADP1 | *A. baylyi* GJS strain with *pcaHG* and *acr1* genes knocked out |
| 4 | *A. baylyi* GJS2_*catA* ADP1 | *A. baylyi* GJS2 strain , with overexpression of *catA* in pBWB162 for efficient bioconversion of catechol to ccMA. |
| 6 | *A. baylyi* GJS2_*aroYkpdBD* ADP1 | *A. baylyi* GJS2 strain with pBWB162_*aroYkpdBD* plasmid, where codon-optimized *aroY* (AB479384) and *kpdBD* genes (AB920346) from *K. pneumoniae* have been cloned into pBWB162 along with the *aroY* gene. |
| 7 | *E.coli_aroY* | Wild-type *aroY* (AB479384) from *K. pneumoniae* cloned into *E.coli* TOP10 |
| 8 | *E.coli_aroYkpdBD* | Codon optimized *aroYkpdBD* from *K. pneumoniae* cloned into *E.coli* TOP10 |
